# The Hippo pathway terminal effector TAZ/WWTR1 mediates oxaliplatin sensitivity in colon cancer cells

**DOI:** 10.1101/2023.03.17.533075

**Authors:** Věra Slaninová, Lisa Heron-Milhavet, Mathilde Robin, Laura Jeanson, Diala Kantar, Diego Tosi, Laurent Bréhélin, Céline Gongora, Alexandre Djiane

## Abstract

YAP and TAZ, the Hippo pathway terminal transcriptional activators, are frequently upregulated in cancers. In tumor cells, they have been mainly associated with increased tumorigenesis controlling different aspects from cell cycle regulation, stemness, or resistance to chemotherapies. In fewer cases, they have also been shown to oppose cancer progression, including by promoting cell death through the action of the P73/YAP transcriptional complex, in particular after chemotherapeutic drug exposure. Using several colorectal cancer cell lines, we show here that oxaliplatin treatment led to a dramatic core Hippo pathway down-regulation and nuclear accumulation of TAZ. We further show that TAZ was required for the increased sensitivity of HCT116 cells to oxaliplatin, an effect that appeared independent of P73, but which required the nuclear relocalization of TAZ. Accordingly, Verteporfin and CA3, two drugs affecting the activity of YAP and TAZ, showed an antagonistic with oxaliplatin in co-treatments. Our results support thus an early action of TAZ to sensitize cells to oxaliplatin, consistent with a model in which nuclear TAZ in the context of DNA damage and P53 activity pushes cells towards apoptosis.

## INTRODUCTION

Colorectal cancer (CRC) is the third leading cause of cancer-related death worldwide (Rawla et al., 2019). 30% of patients present synchronous metastases and 50-60% will develop metastases that will require chemotherapy. The current management of advanced or metastatic CRC is based on fluoropyrimidine (5-FU), oxaliplatin and irinotecan as single agents or more often in combination (e.g. FOLFOX, FOLFIRI, or FOLFIRINOX; Xie et al., 2020). Chemotherapy is combined with targeted therapy including monoclonal antibodies against EGFR (e.g. cetuximab and panitumumab) or VEGF (bevacizumab), tyrosine kinase inhibitors (e.g. regorafenib), and immune checkpoint blockade agents for patients with MSI-High tumors (e.g. pembrolizumab; Xie et al., 2020).

Oxaliplatin is a third-generation platinum antitumor compound with a 1,2-diaminocyclohexane (DACH) ligand (Chaney, 1995; Raymond et al., 1998). It induces mainly intra-strand crosslinks, but also inter-strand crosslinks and DNA-protein crosslinks that stop DNA replication and transcription, leading to apoptotic cell death (Perego and Robert, 2016; Woynarowski et al., 2000, 1998). Oxaliplatin exerts its anti-tumor effect also by inducing immunogenic cell death (Tesniere et al., 2010). Resistance to oxaliplatin can be either intrinsic (primary resistance) or acquired (secondary resistance), and is usually tackled by combining drugs to expose tumoral cells weaknesses or inhibit alternative survival pathways (Vasan et al., 2019). Despite intense research efforts in this field, more information on the molecular mechanisms underlying oxaliplatin mechanism of action are needed to develop new treatment strategies and improve the therapeutic response rate.

The Hippo signaling pathway represents an evolutionarily highly conserved growth control pathway. First discovered through genetic screens in *Drosophila*, it consists of a central cascade of core kinases: MST1/2 and LATS1/2 (homologues of *Drosophila* Hippo and Warts; Heng et al., 2021; Pocaterra et al., 2020; Zheng and Pan, 2019). When activated, LATS1/2 phosphorylate YAP and WWTR1/TAZ (homologues of *Drosophila* Yki), two partly redundant transcriptional co-activators which represent the terminal effector of the Hippo pathway (Reggiani et al., 2021). Phosphorylated YAP and TAZ are retained in the cytoplasm through binding to 14-3-3 proteins, and sent for proteasomal degradation. When the Hippo pathway is not activated, hypo-phosphorylated YAP/TAZ enter the nucleus and bind to specific transcription factors (TFs) to turn on the transcription of target genes. The best characterized TF partners for YAP and TAZ are the TEADs (TEAD1-4, homologues of *Drosophila* Scalloped; Heng et al., 2021; Pocaterra et al., 2020; Zheng and Pan, 2019). While depending on cell type, the classic target genes include genes involved in proliferation, resistance to apoptosis, cytoskeletal remodeling, or stemness (Rosenbluh et al., 2012; Tapon et al., 2002; Totaro et al., 2018). But YAP/TAZ nucleo-cytoplasmic localization (and activity) is also controlled by mechanical cues relayed by the actin cytoskeleton, or by cytoplasmic trapping proteins such as AMOTs (Heng et al., 2021; Pocaterra et al., 2020; Zheng and Pan, 2019). Importantly, the nuclear retention of YAP and TAZ is favored by tyrosine phosphorylation by different kinases, and in particular SRC and YES (Byun et al., 2017; Ege et al., 2018; Li et al., 2016).

The Hippo pathway has been primarily described as a tumor suppressive pathway in a wide variety of solid tumors (Kim and Kim, 2017; Li and Guan, 2022; Nguyen and Yi, 2019; Thompson, 2020; Zheng and Pan, 2019) preventing the pro-tumoral effect of YAP/TAZ. However, in CRCs the role of the Hippo pathway and of YAP and TAZ appears more complex. Several studies point towards a classic pro-tumoral role for YAP and TAZ. In CRC patients tumor samples, high expression and nuclear localizations of YAP correlated strongly with disease evolution and bad prognosis (Ling et al., 2017; Steinhardt et al., 2008), or with resistance to treatments such as 5FU or cetuximab (Kim and Kim, 2017; Lee et al., 2015; Touil et al., 2014). Furthermore, invalidating YAP could blunt tumorigenic behaviors both in mice CRC models (Shao et al., 2014) or in the metastatic HCT116 CRC cell line (Konsavage et al., 2012). However, YAP could exhibit a tumor suppressive role in CRCs. Studies in genetic mouse models have shown that YAP/TAZ restricts canonical Wnt/β-Catenin signaling thus preventing intestinal stem cells amplification, and could act as tumor suppressors in CRCs (Azzolin et al., 2014, 2012; Barry et al., 2013). Similarly, the loss of core Hippo kinases (LATS/MST) was recently shown to inhibit tumor progression in Apc mutant mouse models and in patients-derived xenografts models (Cheung et al., 2020; Li et al., 2020).

The tumor suppressive role of YAP in CRC is further supported by its reported role in response to DNA damage inducer drugs. Studies have shown that, in different cell lines including CRC lines, cell death in response to cisplatin, doxorubicin, or etoposide, is mediated by P73, a protein related to the tumor-suppressor P53. Following treatments, a YAP/P73 complex accumulates in the nucleus, and triggers the transcription of P73 target genes involved in cell death (Lapi et al., 2008; Strano et al., 2005, 2001). The direct interaction between YAP and P73 is proposed to prevent P73 destabilization by the E3-Ubiquitin Ligase ITCH (Levy et al., 2007; Strano et al., 2005). This pro-apoptotic role of YAP is reminiscent to a similar role of Yki in *Drosophila* (a Yki/p53 complex; Di Cara et al., 2015). Importantly, this appears specific to YAP, since TAZ cannot bind to P73, further suggesting that YAP and TAZ, while performing redundant roles, also possess specific activities (Reggiani et al., 2021).

Given that oxaliplatin constitute one of the most used drugs in the treatment of CRCs, it is important to evaluate its effects with respect to the Hippo pathway and to YAP/TAZ which can elicit conflicting roles to oppose or promote CRC tumorigenesis. We show here that upon treatment with oxaliplatin, TAZ accumulated in the nucleus of CRC cell lines. We further show that TAZ was required for early sensitivity of HCT116 to oxaliplatin. Interestingly, the nuclear localisation of TAZ was important, and drugs preventing this such as Dasatinib antagonized the effect of oxaliplatin. These results support an early anti-tumoral role of YAP and TAZ in response to oxaliplatin suggesting particular attention to sequence of treatments and drug combinations should be paid when considering potential future drugging of YAP/TAZ signaling in the treatment of CRCs.

## RESULTS and DISCUSSION

### Oxaliplatin treatment triggers an early cell death program

Oxaliplatin is a third generation platinum compound widely used as part of the first line of treatment for colon cancer patients in the FOLFOX regimen (Chaney, 1995; Raymond et al., 1998; Xie et al., 2020). Inside cells, oxaliplatin binds DNA, generating adducts which ultimately lead to DNA breaks and replicative stress in proliferating cells. When used on proliferating cancer cells, oxaliplatin treatment resulted in concentration-dependent cell death. We measured the IC50 of oxaliplatin on HCT116 colon cancer cells at 0.5µM (Supplemental Figure S1A). This dose reduced the amount of cells by 50% after 4 days of treatment. This dose was about 10 fold lower than the oxaliplatin concentration reported in the blood of treated patients (between 3.7 and 7µM; Graham et al., 2000). The oxaliplatin dose used in this study was thus compatible with the dose that could be ultimately found at the level of tumors in a clinical setting, and did not represent an acute high concentration treatment, highlighting its relevance for studying cellular responses to oxaliplatin.

When treated with oxaliplatin at IC50, HCT116 cells exhibited clear signs of DNA damage such as accumulation of γH2AX, and 53BP1 puncta in the nuclei (Figure 1A&B). Consistently, P53, which has been shown to control a specific cell death program in response to severe DNA damage (for recent reviews Abuetabh et al., 2022; Panatta et al., 2021), accumulated strongly 24h after treatment (Figure 1C). Intriguingly, the P53 related protein P73, previously reported to accumulate and to mediate cell death in response to DNA-damage inducing drugs such as cisplatin, doxorubicin or etoposide, including in HCT116 cells (Lapi et al., 2008; Strano et al., 2005), was destabilized upon oxaliplatin treatment. P63, the third member of the P53 protein family, was not expressed in HCT116, even upon treatment (Figure 1C).

**Figure 1.**
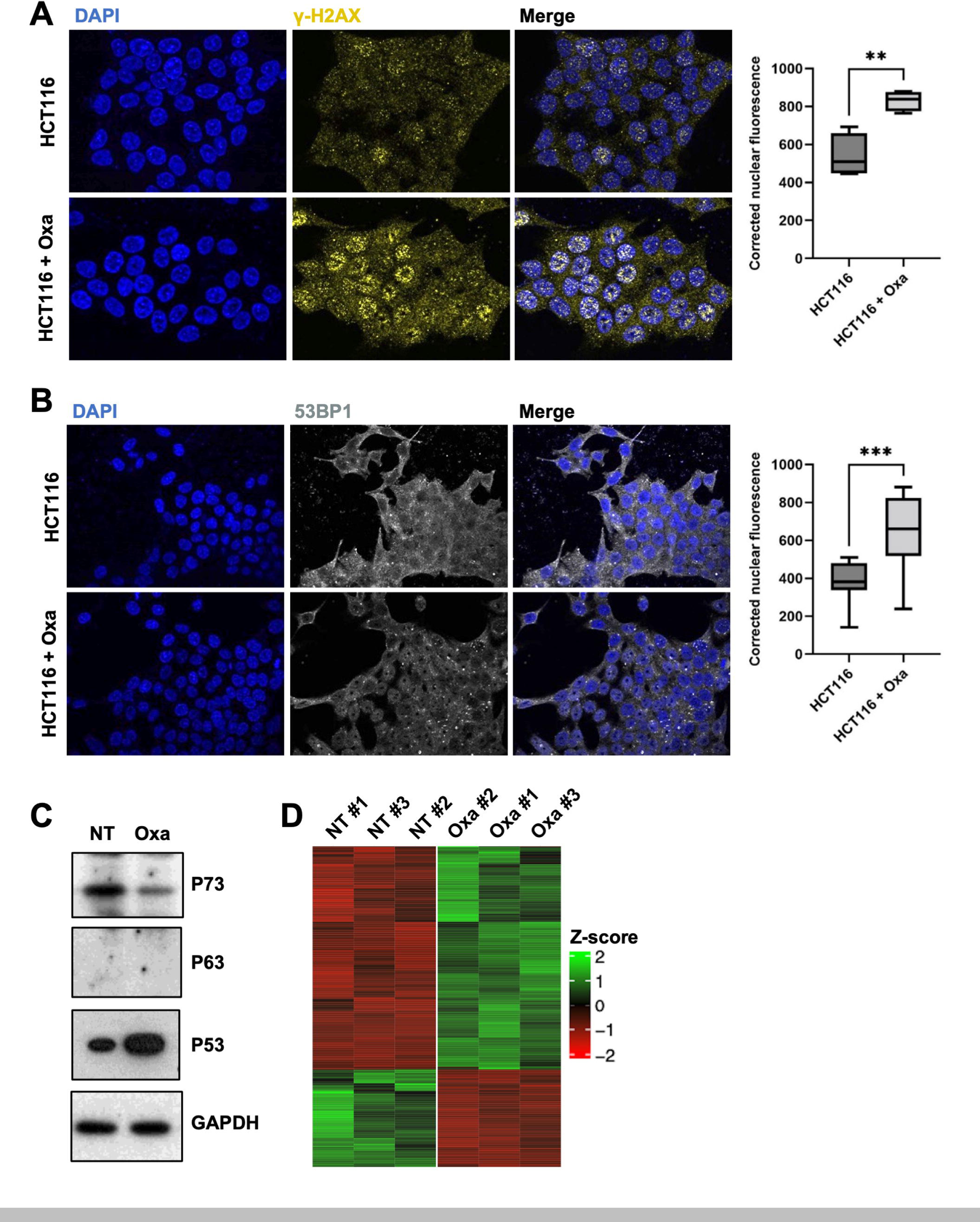
Oxaliplatin treatment induces DNA damage. **A.** Immunofluorescence experiments performed on HCT116 cells treated, or not, with oxaliplatin (0.5 µM) monitoring ψ-H2AX (yellow). DAPI (blue) was used to stain DNA and the nuclei. Quantification of the staining is shown on the right side and is represented as the corrected nuclear fluorescence. Data are represented as the mean ± SEM. (n=3). Unpaired two-tailed Student’s t-test; ** p<0.01. **B.** Immunofluorescence experiments performed on HCT116 cells treated, or not, with oxaliplatin (0.5 µM) monitoring 53BP1 (grey). DAPI (blue) was used to stain DNA and the nuclei. Quantification of the staining is shown on the right side and is represented as the corrected nuclear fluorescence. Data are represented as the mean ± SEM. (n=3). Unpaired two-tailed Student’s t-test; *** p<0.001. **C.** Western blot analysis showing protein expression of TEAD4 and p53 family of proteins in HCT116 treated (Oxa), or not (NT) with oxaliplatin (0.5 µM). GAPDH was used as a loading control (n=3). **D.** Heat map corresponding to the genes differentially expressed in HCT116 cells after 24h of oxaliplatin treatment at IC50 (see Supplemental Table 1). The three replicates for the non-treated (NT) and treated (Oxa) are shown.

To better understand the cellular responses to oxaliplatin we profiled the changes in gene expression after 24h of exposure at IC 50. This analysis revealed that the expression of only a limited number of genes were affected (fold change >1.5, adjusted p-value < 0.05): 253 up-regulated and 111 down-regulated (Figure 1D; Supplemental Table 1). Gene ontology enrichment approaches using the g:profiler online tool (Supplemental Table 2; Raudvere et al., 2019) highlighted that amongst the main cellular processes controlled by the upregulated genes were DNA damage response (GO:0044819 mitotic G1/S transition checkpoint signaling; GO:0000077 DNA damage checkpoint signaling…), apoptosis and cell death (GO:0045569 TRAIL binding; GO:0008219 cell death; GO:0012501 programmed cell death; GO:0006915 apoptotic process …), and p53 response (GO:0072331 signal transduction by p53 class mediator), consistent with the known role of oxaliplatin generating adducts on the DNA. Indeed, many genes up-regulated have previously been associated with p53 signaling, and represent p53 canonical target genes such as CDKN1A/P21, P53I3, BAX, or TIGAR (REAC:R-HSA-3700989 Transcriptional Regulation by P53; WP:WP4963 p53 transcriptional gene network). While up-regulated genes controlled mainly cell death programs, the down-regulated genes were involved in DNA replication (GO:0006260) and cell cycle (GO:0007049) consistent with the well documented effect of DNA damage on blocking cell cycle and proliferation (Abuetabh et al., 2022).

Amongst the genes mis-regulated were also genes related to inflammation and immune cell recruitment (e.g. the upregulated genes CXCR2, EBI3/IL-27, or NLRP1, and the downregulated gene IL17RB) consistent with the previously reported role of oxaliplatin during immune cell death (Tesniere et al., 2010).

Finally, these analyses also highlighted several genes involved in cell architecture, namely cytoskeleton and junctional complexes. Amongst the most striking features were changes in the expression of integrin and extracellular matrix proteins engaging Integrins and Focal Adhesions: collagens COL5A1 and COL12A1, as well as laminins LAMA3, LAMB3, and LAMC1 and integrin ITGA3. These observations suggest that treated cells might remodel their extracellular matrix, their Focal Adhesions, and the signaling pathways associated. The RNA-Seq analyses revealed also many changes to the cytoskeleton, including an upregulation of several keratin-based intermediate filaments (KRT15/19/32) and associated factors (KRTAP2-3 and SFN). Several genes controlling the actin cytoskeleton were also affected such as the branched actin regulators WDR63, CYFIP2, or WASF3, or different genes predicted to control RHO activity (up: RHOD, EZR, and RAP2; down: ARHGAP18).

Taken together, these results suggest that upon oxaliplatin treatment, HCT116 cells implement an early cell death program, which is likely mediated by the elevated P53 levels, and many “bona-fide” P53 direct target genes involved in cell death are upregulated. Unlike other treatments such as cisplatin, doxorubicin, and etoposide (Lapi et al., 2008; Strano et al., 2005), oxaliplatin is unlikely to mobilize the p73 anti-tumoral response since P73 levels are decreased upon oxaliplatin treatment. The difference is striking when considering closely related platinum compounds such as cisplatin and oxaliplatin. This difference is unlikely due to timing as we could not observe any P73 up-regulation after oxaliplatin treatment even after shorter or longer exposures. Even though dose comparisons between different compounds is tricky, we note that the cisplatin dose was 50 times higher than that of oxaliplatin. Alternatively, while both are thought to act primarily as generators of lethal amounts of DNA breaks, their difference in mobilizing either P73 (cisplatin) or P53 (oxaliplatin) might arise from different alternative cellular effects independent of DNA damage.

### Oxaliplatin treatment triggers YAP and TAZ nuclear accumulation

Having established a regimen for treating HCT116 cells with oxaliplatin, and given the complex reported roles of YAP/TAZ in CRCs (see Introduction), we investigated whether YAP/TAZ could be affected, and thus monitored TEAD, YAP, and TAZ expressions and localizations following oxaliplatin treatment.

After 24h (or 48h) of oxaliplatin treatment at the IC50, we did not observe any change in the total levels or in the nuclear localization of TEAD4, the main TEAD paralogue in colon cells (Figure 2A). However, TAZ and YAP nuclear localizations increased following oxaliplatin treatment in our culture conditions: the TAZ and YAP nuclear staining increased by 60% and 55% respectively when compared to untreated cells (Figure 2A). TAZ nuclear accumulation was also observed in two other CRC cell lines: LoVo and Caco-2 (Supplemental Figure S2). TAZ nuclear accumulation was further confirmed by fractionation experiments (Figure 2C; see Materials and Methods). This increase in TAZ nuclear localization was reflected by an increase in total TAZ levels by western blot analysis (Figure 2B). However, YAP total levels, and more importantly the levels of YAP phosphorylation on Serine 127 (S127) were unchanged (Figure 2B).

**Figure 2.**
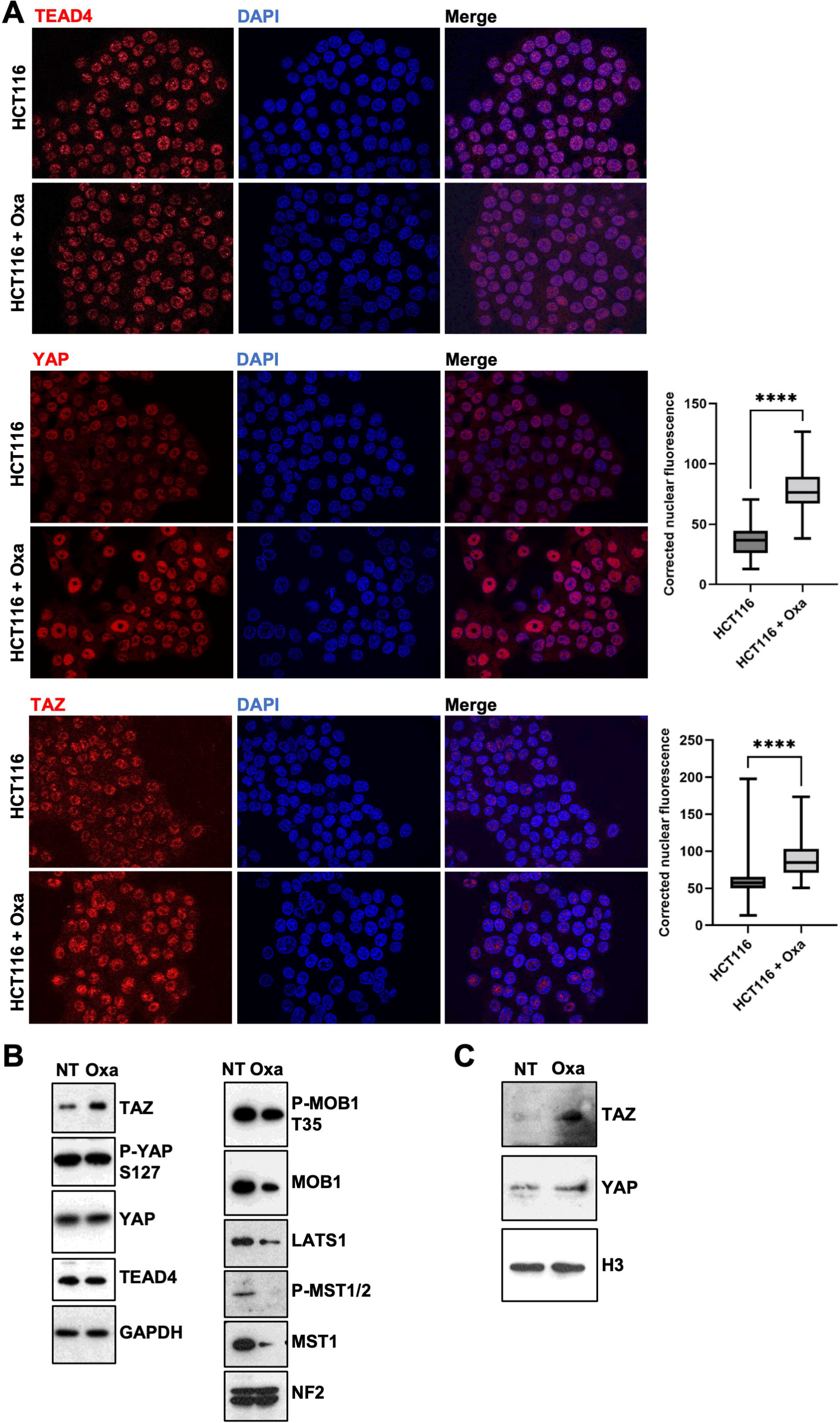
Oxaliplatin treatment triggers YAP and TAZ nuclear accumulation. **A.** Immunofluorescence experiments performed on HCT116 cells treated, or not, with oxaliplatin (0.5 µM) monitoring TEAD4 (top panels), YAP (middle panels), and TAZ (bottom panels) nuclear localization (red). DAPI (blue) was used to stain DNA and the nuclei. Quantification of both stainings are shown on the right side of the figures and are represented as the corrected nuclear fluorescence. Data are represented as the mean ± SEM (n=3). Unpaired two-tailed Student’s t-test; **** p < 0.0001. **B.** Western blot analysis showing protein expression and/or activation of Hippo pathway components in HCT116 cells treated (Oxa), or not (NT) with oxaliplatin (0.5 µM). GAPDH was used as a loading control (n=3). **C.** Western blot analysis after subcellular fractionation showing the relative amount of YAP and TAZ protein in the nuclear fraction of HCT116 cells treated (Oxa), or not (NT) with oxaliplatin (0.5 µM). Histone H3 was used as a nuclear loading control for the fractionation (n=3).

The YAP S127 phosphorylation is deposited by the LATS1/2 Hippo pathway terminal kinases and mediate the cytoplasmic retention of YAP by the 14-3-3 proteins and later targeting for proteasomal degradation (Heng et al., 2021; Pocaterra et al., 2020; Zheng and Pan, 2019). Western-blot analyses on total protein extracts showed that several key proteins in the core Hippo pathway were hypo-phosphorylated (p-MST1/2, p-MOB1) indicating a general lower activity of the core Hippo pathway (Figure 2B). Total protein levels were also lower after treatment, further suggesting a lower Hippo pathway activity in response to oxaliplatin, consistent with the increased TAZ levels and increased nuclear TAZ localization (Figure 2A&C). However, given that levels of phospho-YAP and total YAP remained unchanged, how the Hippo pathway down-regulation could have differing effects on YAP and TAZ remains to be explored. YAP and TAZ appear only partly redundant, and YAP and TAZ specific regulations have been reported (Reggiani et al., 2021). It is noteworthy that an additional phospho-degron is present in TAZ, making it more sensitive to degradation than YAP. This increased sensitivity might magnify TAZ level changes when the Hippo pathway is inhibited by oxaliplatin (Azzolin et al., 2012). The decreased protein levels of different Hippo pathway components in response to oxaliplatin were unlikely due to reduced mRNA abundance, since we did not observe any change in our RNA-Seq, suggesting that it might be a consequence of reduced translation and/or increased protein degradation. Indeed, previous studies have shown that core Hippo pathway components can be regulated by ubiquitination such as LATS1 or MOB1 (Ho et al., 2011; Lignitto et al., 2013; Salah et al., 2011). Whether oxaliplatin treatment triggers a specific ubiquitin-mediated destabilization of the core Hippo pathway remains however to be studied.

### YAP is dispensable for Oxaliplatin-mediated cell death

Performing pathway analyses on the mis-regulated genes highlighted a strong activation of p53 signaling (Supplemental Table S2). Motif enrichment analyses suggested that the p53 family of transcription factors were the main controllers of the up-regulated genes. With the exception of Axl, none of the “classic” YAP/TAZ target genes such as CTGF, CYR61/CCN1, or BIRC2 (or genes involved in cell cycle progression, cytoskeleton regulation, or drug resistance; (Pocaterra et al., 2020; Totaro et al., 2018) were up-regulated after oxaliplatin treatment. We thus wondered what would be the role of YAP and TAZ in the response to oxaliplatin treatment. Indeed, other anti-cancer drugs such as cisplatin have been shown to promote cell death in part through the implementation of a P73/YAP-dependent cell death program. Mechanistically, it has been proposed that DNA damage induced by cisplatin stabilizes YAP which then binds and protects P73 from ITCH-mediated degradation (Levy et al., 2007); the P73/YAP complex accumulates in the nucleus to turn on the expression of P73 target genes involved in cell death (Lapi et al., 2008; Strano et al., 2005, 2001). We thus wondered whether the accumulation of TAZ (and the moderate accumulation of YAP) in the nucleus could also participate in the cell death induced by oxaliplatin.

To test the requirement of YAP and TAZ, we invalidated YAP and TAZ by RNA interference. The sole invalidation of YAP by shRNA led to a very modest reduction in oxaliplatin sensitivity (IC50 in *shYAP* was determined at 0.62 compared to 0.52 in *shLuc* controls) (Figure 3A&C). It is noteworthy that, under the culture conditions used, YAP appeared dispensable for HCT116 cells since the shRNA led to a knock-down efficiency >90%. These results suggest that the cell death in response to oxaliplatin might not be dependent (or only marginally) on the YAP/P73 complex as previously reported for other DNA-damage inducing compounds (Lapi et al., 2008; Levy et al., 2007; Strano et al., 2005), but depends on alternative mechanisms.

**Figure 3.**
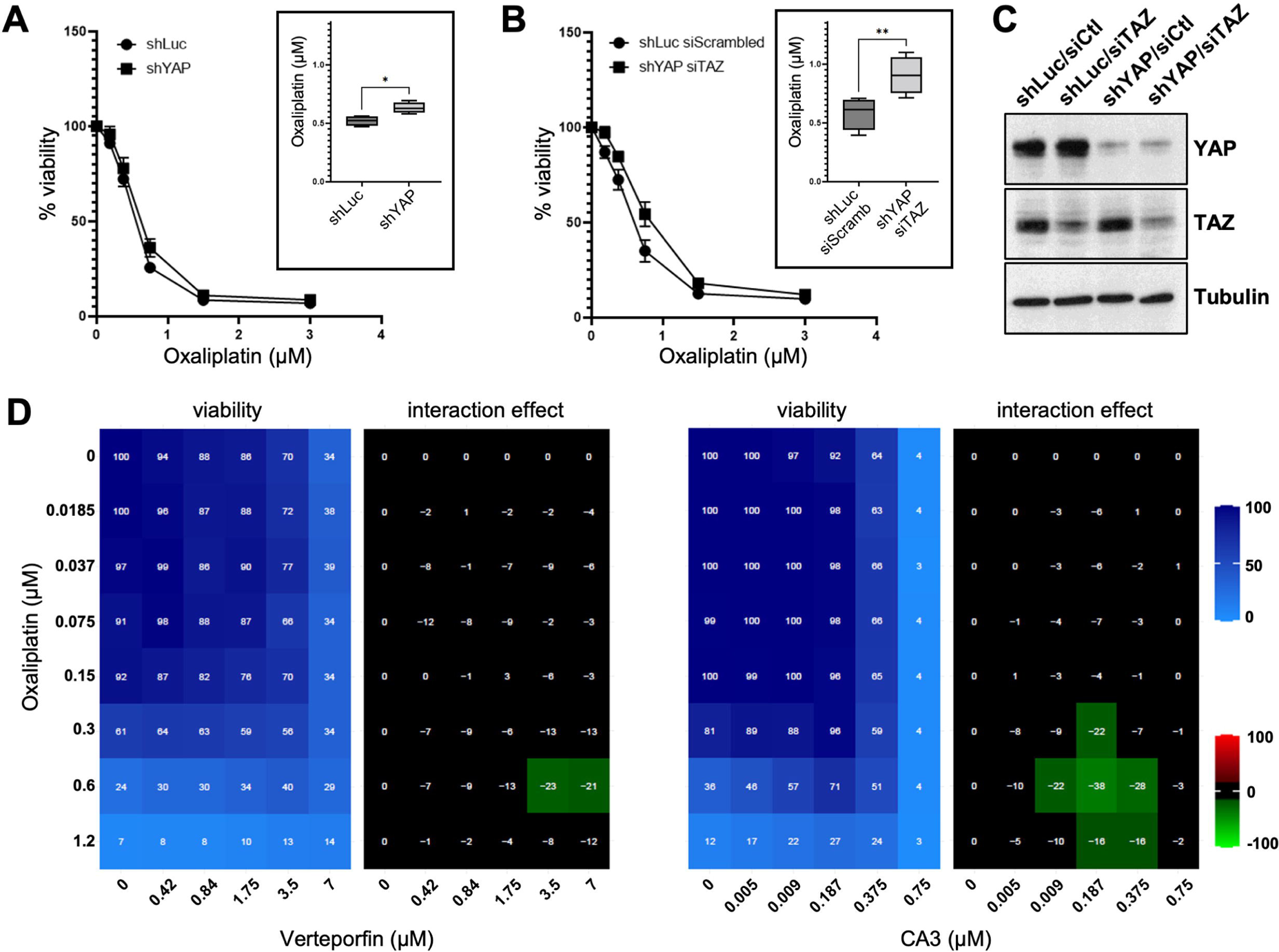
YAP and TAZ are required for oxaliplatin-mediated cell death. **A.** HCT116-*shYAP* and HCT116-*shLuc* cell lines were treated with increasing doses of oxaliplatin for 96h. Cell viability analysis was then assessed using SRB assay and the IC50 of oxaliplatin was calculated as the concentration needed to kill 50% of the cells (shown in the inset). Paired two-tailed Student’s t-test,* p<0.05. **B.** HCT116-*shYAP-siTAZ* and HCT116-*shLuc-siCtl* (control) cell lines were treated with increasing doses of oxaliplatin for 96h. Cell viability analysis was then assessed using SRB assay and the IC50 of oxaliplatin was calculated as the concentration needed to kill 50% of the cells (shown in the inset). Paired two-tailed Student’s t-test, ** p<0.01. **C.** Western blot analysis showing protein expression of TAZ and YAP in HCT116-*shLuc*, *-siTAZ*, *-shYAP* and both *-shYAP-siTAZ* used in panel A and B. Tubulin was used as a loading control (n=3). **D.** HCT116 cells were incubated with increasing concentrations of oxaliplatin and either Verteporfin or CA3. Cell viability was assessed with the SRB assay in 2D to obtain the viability matrix. Drug concentrations were as follows: Verteporfin (from 0.437 to 7 µM), CA3 (from 0.004 to 0.75 µM) and Oxaliplatin (from 0.0185 to 1.2 µM). The synergy matrices were calculated as described in Materials and Methods.

### TAZ promotes cell death in response to oxaliplatin, independently of P73

We then investigated the role of TAZ. Strikingly, while the depletion of YAP had hardly any effect, the combined knock-down of both YAP (*shYAP*) and TAZ (*siTAZ*), resulted in a clear increase in resistance to oxaliplatin, where the IC50 reached 0.91µM in *shYAP/siTAZ* HCT116 cells compared to 0.58µM in *shLuc/siScrambled* HCT116 control cells (Figure 3B&C), highlighting that TAZ participates to cell death in response to oxaliplatin. The effects observed were specific to the *siTAZ*, since we observed a re-sensitization of treated cells when complementing them with an expression vector for a murine version of *Taz* insensitive to the *siTAZ* designed against human *TAZ* (Supplemental Figure S3A&B). We then wondered whether the increased sensitivity promoted by TAZ could be dependent on P73, in a similar mechanism as proposed for cisplatin. However, while P53 accumulated in response to oxaliplatin in HCT116, P73 levels were decreased, undermining the role of P73 in response to this drug (Figure 1C). This absence of P73 stabilization, is consistent with the absence of increased YAP levels after oxaliplatin treatment (Figure 2B). These results highlight that, although overexpressed YAP could bind and stabilize endogenous P73 (Supplemental Figure S4; Levy et al., 2007; Strano et al., 2005, 2001), oxaliplatin treatments at the clinically relevant doses used, do not lead to YAP and P73 stabilization. We then confirmed that TAZ cannot bind P73 (Supplemental Figure S4), ruling out that the elevated nuclear TAZ following oxaliplatin could act through a transcriptional complex with P73 to enhance cell death. A recent study reported a direct interaction between TAZ and P53 in MCF7 and HCT116 cells, which resulted in the inhibition of P53 activity towards senescence (Miyajima et al., 2020). However, when we performed co-immunoprecipitation experiments, we were unable to document any interaction between over-expressed YAP or overexpressed TAZ with endogenous P53 in normal or oxaliplatin treated HCT116 cells (Supplemental Figure S4). Furthermore, the increased oxaliplatin resistance of cells upon *YAP/TAZ* knockdown supports strongly that TAZ acts to promote cell death and thus cooperates with P53 rather than antagonizes its activity as suggested before (Miyajima et al., 2020). Taken together, these results suggest that the sensitivity of HCT116 cells to oxaliplatin mediated by YAP and TAZ is not mediated by the direct interaction of YAP or TAZ to P53 or P73.

### Increased resistance to oxaliplatin upon YAP/TAZ activity blockade

The sh/siRNA interference results suggested that TAZ was required for sensitivity to oxaliplatin. To validate independently the knock-down experiments, we used a pharmacological approach with drugs targeting YAP/TAZ activity and monitored their action in combination with oxaliplatin. We performed 2D matrices co-treatment analyses in which cells were treated with increasing amounts of oxaliplatin and of the YAP/TAZ inhibitors verteporfin or CA3 (Figure 3D and Supplemental Figure S1B &C; Liu-Chittenden et al., 2012; Song et al., 2018). In both cases, the co-treatments led to a marked increase in the HCT116 resistance to oxaliplatin. Similar results were obtained on two other CRC cell lines: LoVo, and Caco-2 (Supplemental Figure S3C&D). The mode of action of verteporfin remains unclear and might involve increased retention in the cytoplasm of YAP and TAZ, or their degradation, preventing them from complexing in the nucleus with their transcription factor partners (Wang et al., 2016). A recent study showed that CA3 reduced the transcriptional activity mediated by YAP/TAZ-TEAD (reduction in target genes expression), with only minor effects on YAP protein levels (Morice et al., 2020). Even though the exact mode of action of verteporfin and CA3 remain unclear, the increased resistance to oxaliplatin observed by co-treating cells with YAP/TAZ pharmacological inhibitors, confirms the results obtained with the genetic knock-down, and supports a model where increased TAZ activity participate in the sensitivity of CRC cells to oxaliplatin.

### Src inhibition by Dasatinib reduces HCT116 cells sensitivity to oxaliplatin

The results suggest thus that preventing TAZ signaling in the early phases of oxaliplatin treatment would represent a counter-productive approach, leading to reduced efficacy of oxaliplatin to induce cell death. Besides the canonical Hippo signaling pathway, the nucleo-cytoplasmic shuttling of YAP and TAZ is under the control of many other inputs. In particular, YAP and TAZ retention in the nucleus is promoted by the action of different tyrosine kinases, such as ABL or SFKs (Src Family Kinases) which phosphorylate the C-termini of YAP and TAZ (Y357 or Y316 respectively; Byun et al., 2017; Ege et al., 2018; Guégan et al., 2022; Kedan et al., 2018; Lamar et al., 2019; Li et al., 2016). Due to its high relevance for colon cancer, we focused our analysis on SRC, frequently activated in colon carcinoma (Sirvent et al., 2020). An earlier study showed that depending on the colon cancer cell line considered, SRC could be activated, inhibited, or not affected following oxaliplatin treatment (Kopetz et al., 2009). We could replicate that SRC was not activated after 24h of oxaliplatin treatment in HCT116 cells (as measured by phosphorylation on Y416; Figure 4A). Working with HCT116, we are thus in a position to test the contribution of SRC to YAP/TAZ shuttling during oxaliplatin treatment without the complications arising from treatment-induced acute SRC activation. Previous reports suggested that the classic SRC kinase inhibitor Dasatinib could be used as a drug to prevent YAP/TAZ signaling (Rosenbluh et al., 2012). Indeed, combining Dasatinib with oxaliplatin treatment, prevented the nuclear accumulation of TAZ (Figure 4B). The addition of Dasatinib to oxaliplatin treated cells led to a dramatic reduction of the TAZ nuclear staining when compared to oxaliplatin alone (95% reduction; see Materials and Methods). It should be noted however, that Dasatinib treatment at 50nM reduced slightly the elevated global TAZ levels observed in response to oxaliplatin (Figure 4A). Nevertheless, even though TAZ appeared a bit more unstable in presence of Dasatinib, its nucleo-cytoplasmic ratio was still profoundly affected by Dasatinib, preventing nuclear accumulation (Figure 4B).

**Figure 4.**
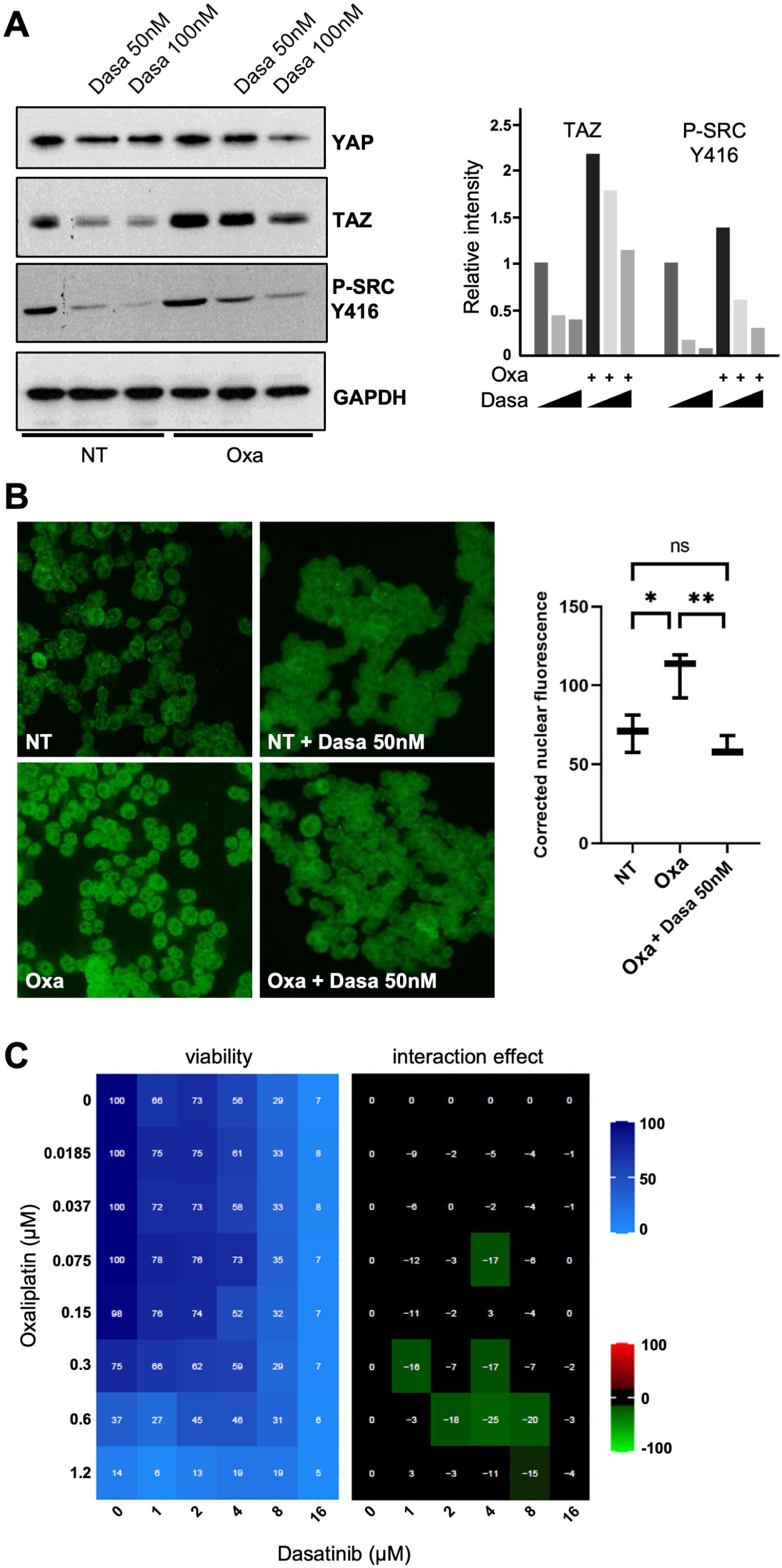
Src inhibition by Dasatinib reduces HCT116 cells sensitivity to oxaliplatin. **A.** Western blot analysis showing protein expression of YAP and TAZ in HCT116 treated, or not, with oxaliplatin (0.5 µM) and/or Dasatinib (50 nM and 100 nM). Phopsho-SRC blotting was used to evaluate the inhibition of SRC activity using Dasatinib. GAPDH was used as a loading control (n=3). Quantification of the blots (performed using Image J software) is shown on the right side of the figure. **B.** Immunofluorescence experiments performed in HCT116 cells treated, or not, with oxaliplatin (0.5 µM) and/or Dasatinib (50 nM) monitoring YAP (top panels) and TAZ (bottom panels) nuclear localization (red). DAPI (blue) was used to stain DNA and the nuclei. Quantification of both stainings are shown on the right side of the figures and are represented as the corrected nuclear fluorescence. Data are represented as the mean ± SEM (n=3). Unpaired two-tailed Student’s t-test; ****p < 0.0001. **C.** HCT116 colorectal cancer cell lines were incubated with increasing concentrations of oxaliplatin and Dasatinib. Cell viability was assessed with the SRB assay in 2D to obtain the viability matrix. Drug concentrations were as follows: Dasatinib (from 1 to 16 µM) and oxaliplatin (from 0.0185 to 1.2 µM). The synergy matrix was calculated as described in Materials and Methods.

We thus asked what would be the combined effect of Dasatinib treatment and oxaliplatin in HCT116 cells. We thus performed 2D matrices co-treatment analyses in which cells were treated with increasing amounts of oxaliplatin and of Dasatinib using drug ranges encompassing their respective IC50 (0.5µM for oxaliplatin and 8µM for Dasatinib; Supplemental Figure S1A&D). Strikingly combining both drugs showed clear regions of antagonism, suggesting that Dasatinib treatment reduced HCT116 cells sensitivity to oxaliplatin (Figure 4C). These results further support a model in which the nuclear relocalization of TAZ in response to oxaliplatin treatment sensitizes cells, and caution the use of Dasatinib in combination to oxaliplatin.

### YAP/TAZ promote cell death in the early response to chemotherapeutic agents

Taken together the results presented here show that oxaliplatin promotes the fast nuclear relocalization of TAZ which then participates to the cells sensitivity to oxaliplatin. Given that we could not find any interaction between TAZ and P53 family members, but that the nuclear localization of TAZ is required for its effect, we could envision several models:

i. either the TAZ/TEAD transcription complex, in the context of DNA damage and P53 activation, promotes the transcription of specific early response genes promoting cell death;
ii. or the slight increase at the transcriptional level of “classic” YAP/TAZ/TEAD targets involved in proliferation sensitizes cells to DNA damage and replicative stress;
iii. or alternatively, TAZ acts through a new complex involved in cell death, independently of TEAD.

More studies should help to distinguish between these potential models.

YAP and TAZ, have been implicated in the resistance to various chemotherapies or targeted therapies in different cancers (Kim and Kim, 2017; Nguyen and Yi, 2019; Zeng and Dong, 2021). It should be noted that the current study focuses on the immediate effects of oxaliplatin within the first hours after exposure. Whether YAP and TAZ are later important for the maintenance of the resistance acquired by the surviving clones is not addressed in this study. Hints towards this later role of YAP/TAZ, are suggested by the elevated YAP levels reported in many cancer cells following resistant clone selection (our own unpublished results, and (Kim and Kim, 2017; Nguyen and Yi, 2019; Zeng and Dong, 2021). Functional studies impairing YAP demonstrated that YAP is indeed required for the tumorigenicity of resistant cells (Yoshikawa et al., 2015). Furthermore, elevated YAP and TAZ nuclear staining is frequently observed in patients tumor samples, including in CRCs (Li and Guan, 2022; Ling et al., 2017; Steinhardt et al., 2008; Thompson, 2020). In advanced cancers, almost all patients undergo one or more rounds of treatment before surgery, if surgery is possible. It is thus unclear whether the increased YAP/TAZ nuclear levels observed in tumor samples reflect primary response to treatment (as suggested by the current study), or whether they represent a secondary state that might have been selected in the cells resistant to treatment.

The current study investigates the early response to oxaliplatin, supporting an early tumor suppressive role of YAP/TAZ in response to treatment, in which, in the context of detrimental DNA damage, YAP/TAZ activity promotes cell death. Is this role general or is it specific to CRCs and oxaliplatin? Independently of the mechanism involved (YAP/P73 complex as previously reported or alternative TAZ-mediated mechanisms as shown here), different breast and colon cancer cell lines mobilize YAP or TAZ to promote cell death in response to many different DNA damaging agents (Basu et al., 2003; Lapi et al., 2008; Levy et al., 2007; Strano et al., 2005, 2001). This anti-tumoral role appears evolutionarily conserved and in *Drosophila* the YAP/TAZ homologue Yki promotes cell death in response to different stress inducing agents (Di Cara et al., 2015), further suggesting that YAP/TAZ might promote cell death in response to chemotherapeutic agents in other cancers beside CRCs and breast cancers. When considering drugging YAP/TAZ signaling in the treatment of CRCs and other cancers, special attention should thus be given to drug combinations, and importantly the sequence in which they will be used.

## MATERIALS AND METHODS

### shRNA construction

shRNA directed against human *YAP*, *TAZ*, or the non-relevant *Luciferase* gene were designed by adding to the selected targeted sequences, overhangs corresponding to BamHI and EcoRI cloning sites at the 5’end of forward and reverse strand, respectively. Resulting oligos were then annealed together and cloned into the pSIREN-RetroQ vector (TaKaRa) according to the manufacturer’s protocol between BamHI and EcoRI cloning sites.

Targeted sequences:

*shRNA-YAP(3619):* CAATCACTGTGTTGTATAT

*shRNA-TAZ(1417):* CCCTTTCTAACCTGGCTGT

*shRNA-Luciferase:* CGTACGCGGAATACTTCGA

### Cell culture and cell transfections

Certified Human HCT 116 and LoVo colorectal cancer cell lines (RRID:CVCL_0291, RRID:CVCL_0399) were obtained from LGC Standards (ATCC-CCL-247, ATCC-CCL-229). Caco-2 cells (RRID:CVCL_0025) were certified independently. Cells were cultured in RPMI1640 supplemented with 10% FBS at 37°C in a humidified atmosphere with 5% CO_2_. Cultures were regularly checked to be mycoplasma-free. No antibiotics were used to avoid any cross-reaction with the Oxaliplatin treatment.

HCT116 cells expressing shRNA against *YAP*, *TAZ*, or *Luciferase* (Luc; control) were obtained by retroviral gene transduction of the corresponding pSIREN vectors. Retroviral particles were produced in HEK293 cells and subsequently used to infect HCT116 cells. Positive clones were selected with 1 μg/mL puromycin and pooled together.

HCT116-*shYAP/siTAZ* cells were created by transfecting 100nM of *TAZ siRNA* (Dharmacon siGENOME SMARTpool *#*M-016083-00-0005) into HCT116-*shYAP* cells using Lipofectamine 2000 (Invitrogen) according to manufacturer’s protocol. As a negative control, 100nM of *siScrambled* (D-001206-13) was transfected into HCT116-*shLuc* cells.

Murine Taz was expressed by transfecting cells with pEF-TAZ-N-Flag from Michael Yaffe (Addgene #19025; RRID:Addgene_19025; Kanai et al., 2000).

### RNA-Seq

HCT 116 cells were plated to reach 60 to 70% of confluence and treated with 0.5 μM Oxaliplatin (IC_50_) for 24 hours. RNA was extracted using RNeasy plus mini kit (Qiagen), quantified and analyzed for its integrity number (RIN) using a Bioanalyzer (Agilent 2100 at the IRMB: https://irmb-montpellier.fr/single-service/transcriptome-ngs/). RNA (1 µg) with RIN between 8 and 10 were sent for RNA-Sequencing analysis to Fasteris biotechnology company (http://www.fasteris.com). After library preparation, sequencing was performed on the Illumina NovaSeq 6000 platform (S1 2×100 full FC). Mapping on the human genome GRCh38 was performed using the protocol STAR 2.7.5b leading to 80-100 Millions reads per condition. Normalization and pairwise differential expression analyses were performed using the R package DESeq2 (2.13) (Anders and Huber, 2010).

RNA-Seq data has been deposited in NCBI’s Gene Expression Omnibus under accession number GSE227315.

### Western blotting

Proteins issued from transfected untreated and treated HCT 116 cells were extracted, analyzed by SDS-PAGE. Dilutions and antibodies’ references are listed below.

Flag M2 (1/2000; Sigma-Aldrich #F1804)

GAPDH (1/3000; Proteintech #60004)

Histone H3 (1/1000; Cell Signaling Technology CST #4499)

LATS1 (1/1000; CST #3477)

MOB1 (1/1000; CST #13730)

p-MOB1 (1/1000; CST #8699)

MST1 (1/1000; CST #3682)

p-MST1/2 (1/1000; CST #49332)

NF2 antibody (Proteintech #26686-1-AP)

P53 (1/5000; Proteintech #10442-1-AP)

P63 (1/250; Santa Cruz #sc25268)

P73 (1/1000; CST #14620)

p-SRC Y416(1/1000; CST #2105)

TAZ (1/1000; CST #4883)

TEAD4 (1/250; Santa Cruz #sc101184)

Tubulin (1/10000; Sigma-Aldrich #T6074)

YAP (1/1000; CST #14074)

p-YAP S127 (1/1000; CST #13008)

### Immunoprecipitation and co-immunoprecipitation

Protein extracts were prepared in lysis buffer (NaCl 150 mM, Tris pH 7.4 10 mM, EDTA 1 mM, Triton X-100 1%, NP-40 0.5%, cOmplete, EDTA-free protease inhibitors (Roche #11873580001) for 30 min on ice before centrifugation. Immunoprecipitations were performed overnight at 4°C on a rocking wheel using mouse EZview Red anti-Flag M2 affinity gel (Sigma-Aldrich #F1804) after transfections of either p2xFlag CMV2 (empty vector), p2xFlag CMV2-YAP2 (YAP1; Addgene #19045) or p2xFlag CMV2-WWTR1 (TAZ). After Flag immunoprecipitation, washes in lysis buffer were performed, followed by protein elution by competition with 3XFLAG peptide (150 ng/µL final concentration) during 1 hour at 4°C. The different immunoprecipitates were then subjected to Western blotting for detection of protein complexes.

### Immunofluorescence

Cells seeded on glass coverslips were fixed 10 min in paraformaldehyde (4 %), before being permeabilized in PBS / 0.1% TritonX-100 for 10 min. After blocking in PBS / 0.5% BSA, cells were incubated with primary antibodies overnight at 4C. Primary antibodies used are listed below. Secondary Alexa Fluor Antibodies (1/600; Invitrogen) were used as described previously (Kantar et al., 2021) for 1 hour at room temperature before mounting the coverslips with Vectashield (Vector Laboratories #H-1200) and imaging on Zeiss Apotome or Leica Thunder microscopes.

Antibodies used were rabbit anti-53BP1 (1/100; CST #4937), mouse anti-phospho-Histone H2AX clone JBW301 (1/200; Millipore #05-636), anti-TAZ (1/100; CST #4883), mouse anti-TEAD4 (1/50; Santa Cruz #sc101184), and rabbit anti-YAP (1/100; CST #14074).

### Nuclear staining quantifications in HCT116 cells

Quantification was performed using ImageJ. Binary mask corresponding to the cell nuclei was based on DAPI staining. Two nuclei touching each other (and therefore recognized as one on binary mask) were manually separated by drawing a 2-pixel line between them. All incomplete nuclei on the edge of the image as well as those that were in mitosis or mechanically damaged were excluded from the analysis. The total signal was calculated as “corrected total cell fluorescence” (CTCF) according to the following formula:

*CTCF = Integrated Density – (Area of selected cell * Mean fluorescence of the background)*

Background fluorescence was measured on three different spots (roughly the size of cell nucleus) outside of the cell. In case of 53BP1 and g-H2AX staining, the whole area covered by the nuclear mask was quantified as one. For YAP and TAZ nuclear staining, each cell was quantified separately using particle analysis tool. Cytoplasmic levels of YAP and TAZ were not quantified due to the small size of the cytoplasm in HCT116 cells.

### IC50 calculation and cytotoxicity

Cell growth inhibition and cell viability after incubation with Oxaliplatin (Sigma Aldrich #O9512), Verteporfin (Sigma Aldrich #SML0534), CA3 (CIL56, Selleckchem, #S8661) or Dasatinib (Selleckchem #S1021) were assessed using the sulforhodamine B (SRB) assay. Exponentially growing cells (750 cells/well) were seeded in 96-well plates in RPMI-1640 medium supplemented with 10% FCS. After 24 hours, serial dilutions of the tested drugs were added and each concentration was tested in triplicate. After 96 hours, cells were fixed with 10% trichloroacetic acid and stained with 0.4% SRB in 1% acetic acid. SRB-fixed cells were dissolved in 10 mmol/L Tris–HCl and absorbance at 540 nm was read using an MRX plate reader (Dynex, Inc., Vienna, VA, USA). IC50 was determined graphically from the cytotoxicity curves.

For HCT116-*shYAP/siTAZ*, cells were transfected in 6 well plates 24h before starting the cell growth and cytotoxicity assays.

### Quantification of the interaction effect

The interaction between the drugs tested *in vitro* was investigated with a concentration matrix test, in which increasing concentration of each single drug were assessed with all possible combinations of the other drugs. For each combination, the percentage of expected growing cells in the case of effect independence was calculated according to the Bliss equation (Greco et al., 1995):

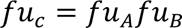

where *fu*_c_ is the expected fraction of cells unaffected by the drug combination in the case of effect independence, and *fu*_A_ and *fu*_B_ are the fractions of cells unaffected by treatment *A* and *B,* respectively. The difference between the *fu*_c_ value and the fraction of living cells in the cytotoxicity test was considered as an estimation of the interaction effect, with positive values indicating synergism and negative values antagonism.

## ACKNOWLEDGEMENTS

The authors thank the different lab members of the Djiane and Gongora teams for helpful discussions. VS was supported by Fondation de France. MR was supported by LabEx NUMEV. DK was supported by Ligue Contre le Cancer. This work was supported by grants from Fondation ARC (#PJA 20141201630) and Ligue Nationale Contre le Cancer (Région Languedoc-Roussillon) to LHM and AD. Work in the lab of CG was also supported by grants and funds from INSERM, the Institut du cancer de Montpellier (ICM), SIRIC (SIRIC Montpellier Cancer Grant «INCa-DGOS-Inserm 6045»), Cancéropole GSO, and the program «investissement d’avenir» (grant agreement: Labex MabImprove, ANR-10-LABX-53-01).

## AUTHOR CONTRIBUTIONS

VS, LHM, LJ, and DK performed experiments. CG and AD designed the experiments. VS, LHM, MR, DK, CG and AD analyzed the data. VS, LHM, DT, LB, CG, and AD interpreted the data. CG and AD wrote the manuscript.

## SUPPLEMENTAL FIGURE LEGENDS

**Supplemental Figure S1. Cell viability in response to drugs**

Cells were treated with increasing doses of oxaliplatin (**A**), Verteporfin (**B**), CA3 (**C**) and Dasatinib (**D**) for 96h. Cell viability analysis was then assessed using SRB assay and the IC50 of each drug could be calculated as the concentration that reduced cell numbers by 50% (shown in the insets).

**Supplemental Figure S2. TAZ nuclear accumulation in LoVo and Caco-2 CRC cell lines**

**A-B**. Immunofluorescence experiments performed on LoVo (A) and Caco-2 (B) cells treated or not, with oxaliplatin at IC50 (0.6 μM and 0.3 μM respectively) monitoring TAZ nuclear localization (green). DAPI (blue) was used to stain DNA and the nuclei. Quantifications are shown on the right side of the figure and are represented as the corrected nuclear fluorescence. Data are represented as the mean ± SEM. Unpaired two-tailed Student’s t-test; **** p < 0.0001.

**Supplemental Figure S3. TAZ mediates sensitivity to oxaliplatin in CRC cell lines**

**A.** HCT116-*shYAP-siScramb* (control), HCT116-*shYAP-siTAZ,* and HCT116-*shYAP-siTAZ* transfected with a Flag tagged murine Taz *(pEFmTaz)* cell lines were treated with increasing doses of oxaliplatin for 96h. Cell viability analysis was then assessed using SRB assay and the IC50 of oxaliplatin was calculated as the concentration needed to kill 50% of the cells. Paired two-tailed Student’s t-test, ** p<0.01, ns non-significant.

**B.** Western blot analysis showing protein expression of TAZ and Flag in the different cell lines used in panel A. GAPDH was used as a loading control.

**C-D**. LoVo (C) and Caco-2 (D) cells were incubated with increasing concentrations of oxaliplatin and either Verteporfin or CA3. Cell viability was assessed with the SRB assay in 2D to obtain the viability matrix. Drug concentrations were as follows: Verteporfin (from 0.875 to 14 µM), CA3 (from 0.15 to 2.4 µM) and Oxaliplatin (from 0.075 to 4.8 µM). The synergy matrices were calculated as described in Materials and Methods.

**Supplemental Figure S4. Interaction between YAP/TAZ and P53 protein family members in HCT116 cells**

Western blot analysis of Flag (YAP or TAZ) immunoprecipitates on protein extracts from HCT116 treated (Oxa), or not (NT) showing an interaction between YAP and p73 (n=3).

**Supplemental Table 1. Differentially expressed genes in HCT116 cells after oxaliplatin treatment for 24h at IC50**

Only differentially expressed genes with an adjusted p-value< 0.05 are shown. Columns are: test_id: position and description of the feature tested

ENSG_ID: Ensemble gene ID

symbol: gene symbol

sample_1: first group in the comparison; untreated cells

sample_2: second group in the comparison; cells treated with oxaliplatin

mean_1: mean of normalized count for the first group in the comparison

mean_2: mean of normalized count for the second group in the comparison

log2FoldChange: log2 fold change estimates

pvalue: pvalue

padj: pvalue adjusted for multiple testing with the Benjamini-Hochberg procedure, which controls false discovery rate (FDR)

**Supplemental Table 2. g:profiler functional enrichment analyses**

Columns are :

GO_ID: Gene Ontology

KEGG_ID: KEGG pathways

REAC_ID: Reactome pathways

WP_ID: WikiPathways

TF_ID: regulatory motif matches from TRANSFAC

MIRNA_ID: miRNA targets from miRTarBase

CORUM_ID: protein complexes from CORUM

HPA_ID: tissue specificity from Human Protein Atlas

HP_ID: human disease phenotypes from Human Phenotype Ontology

Description: description of the functional group

p.Val: p-value

FDR: false discovery rate

Genes: genes found in the intersection

**Supplemental Table 3. RNA-Seq statistics**

