## Supplementary figures and images for "The Hippo pathway terminal effector TAZ/WWTR1 mediates oxaliplatin sensitivity in colon cancer cells"

### Supplemental Figure S1

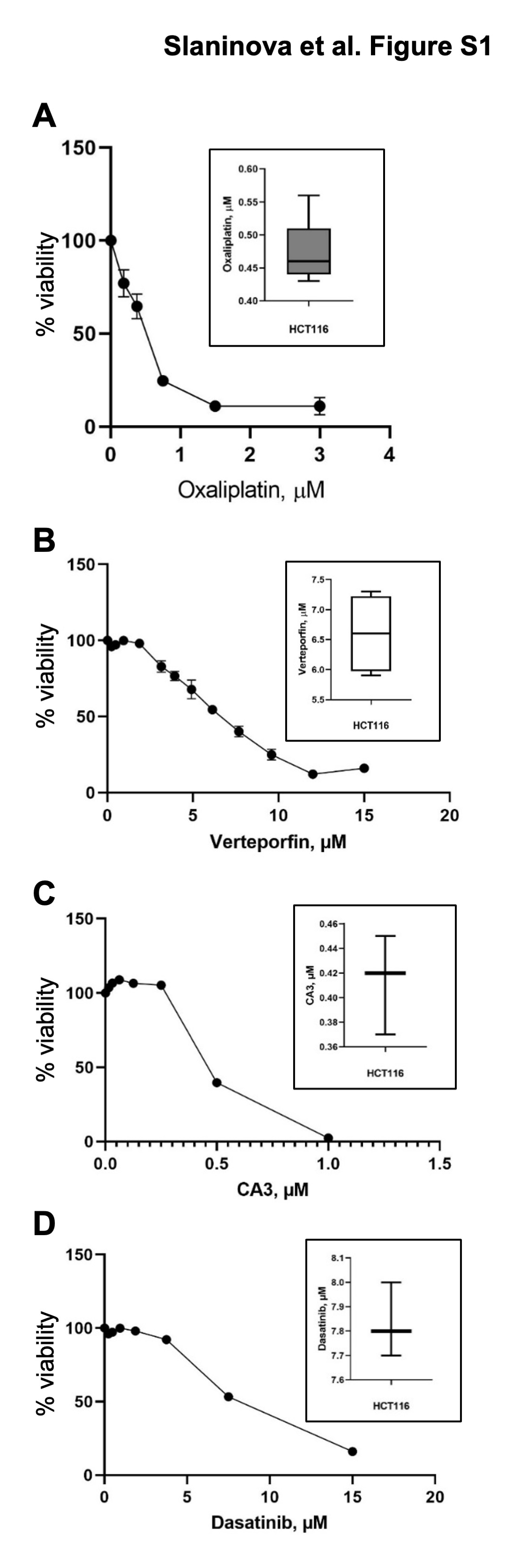

### Supplemental Figure S2

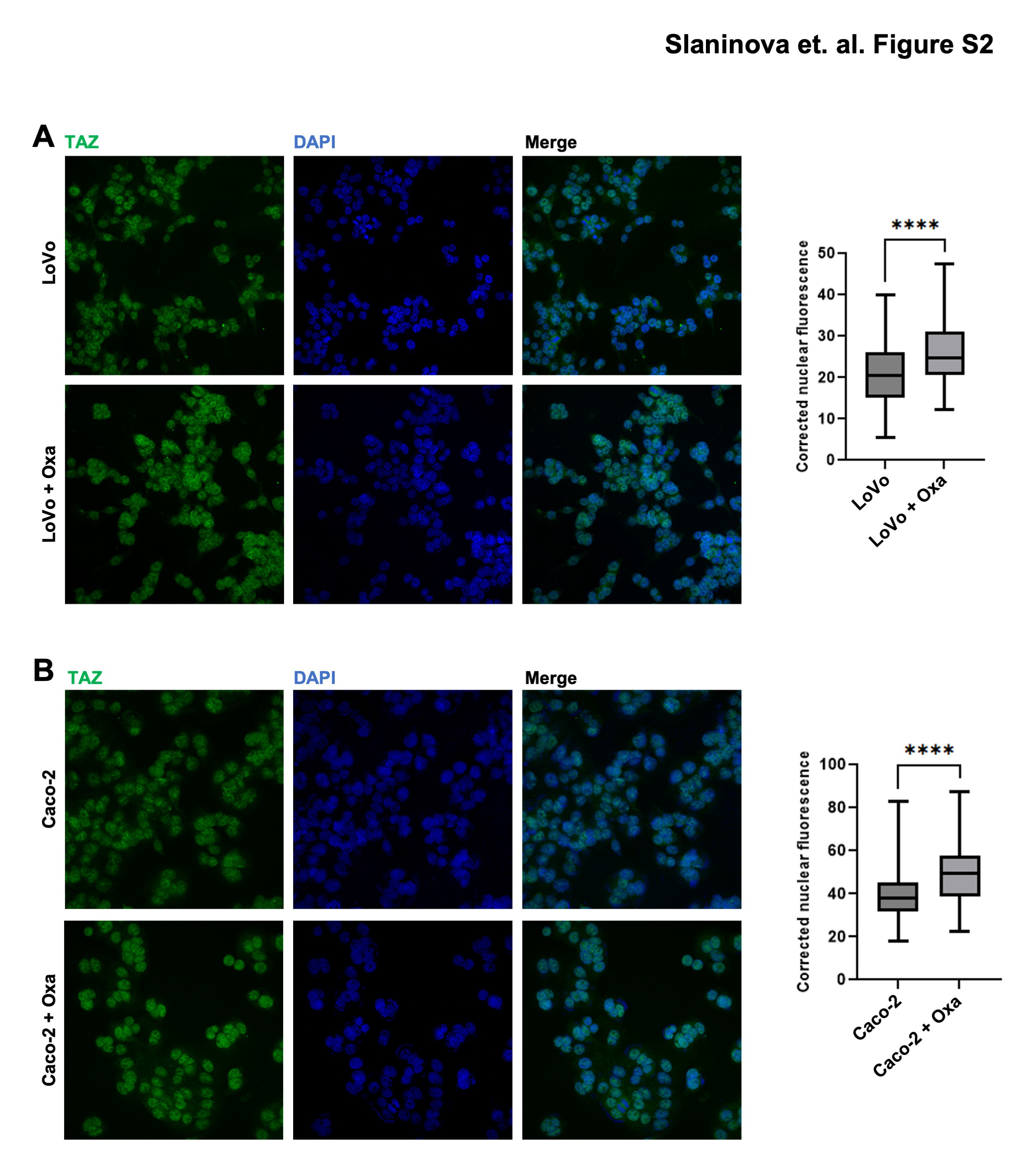

### Supplemental Figure S3

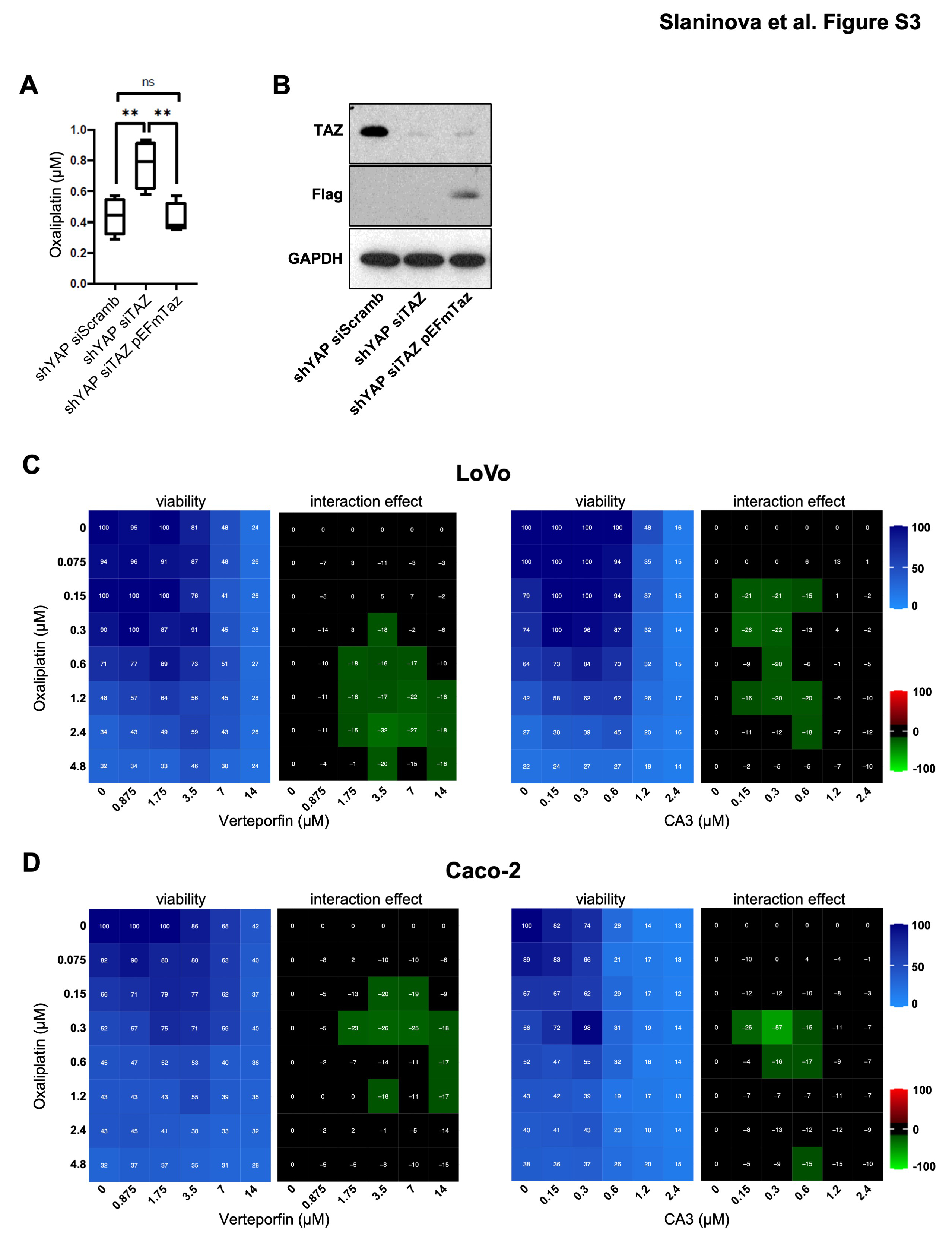

### Supplemental Figure S4

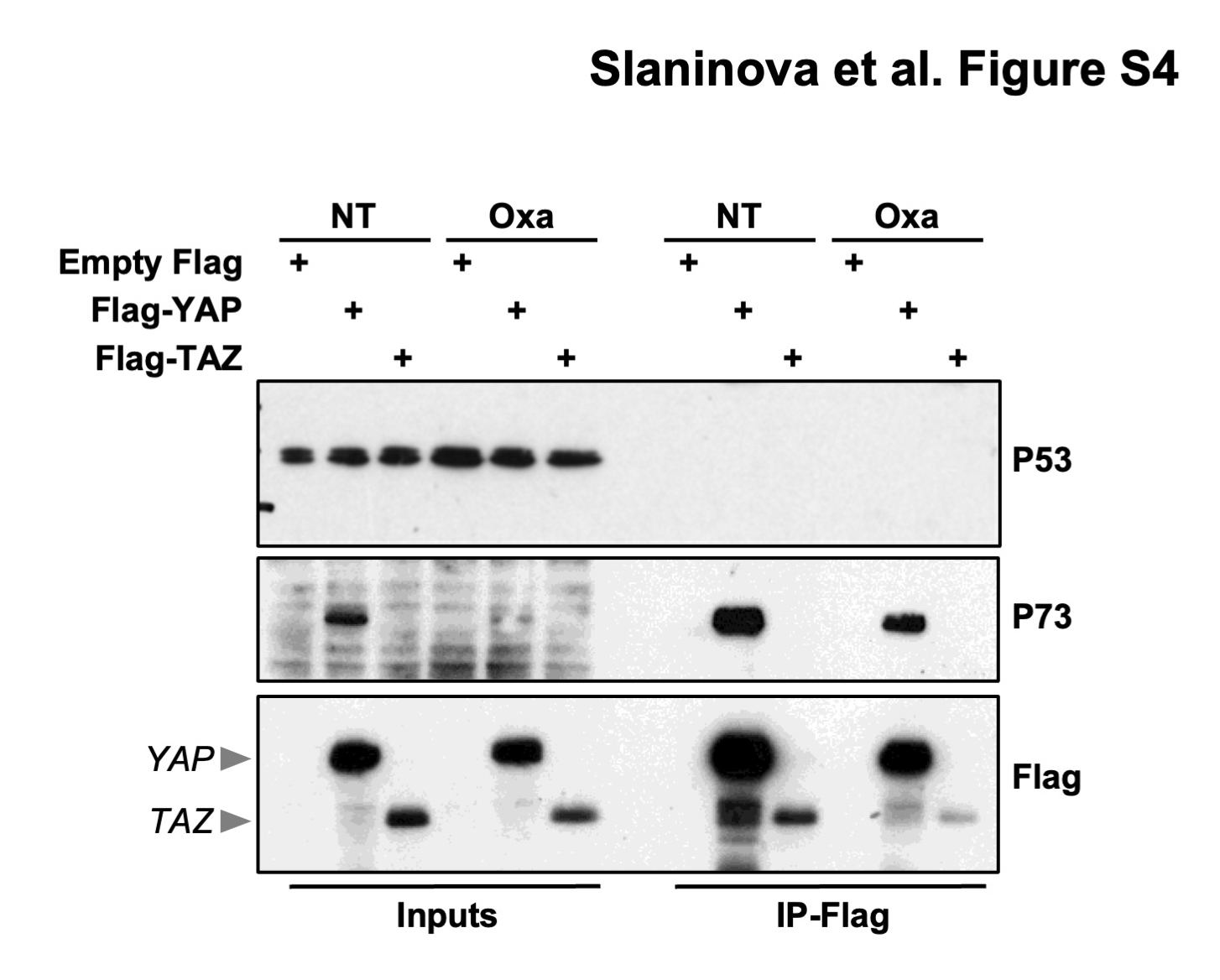
